# Revisiting degron motifs in human AURKA required for its targeting by APC/C-FZR1

**DOI:** 10.1101/2022.01.31.478464

**Authors:** Ahmed Abdelbaki, Camilla Ascanelli, Cynthia N. Okoye, Begum Akman, Giacomo Janson, Mingwei Min, Chiara Marcozzi, Anja Hagting, Rhys Grant, Maria De Luca, Italia Anna Asteriti, Giulia Guarguaglini, Alessandro Paiardini, Catherine Lindon

## Abstract

Mitotic kinase Aurora A (AURKA) diverges from other kinases in its multiple active conformations that may explain its interphase roles and association with cancer, and the limited efficacy of drugs targeting the kinase pocket. Regulation of AURKA activity by the cell is critically dependent on destruction mediated by the Anaphase-Promoting Complex (APC/C^FZR1^) during mitotic exit and G1 phase and requires an atypical N-terminal degron in AURKA called the ‘A-box’ in addition to a reported canonical D-box degron in the C-terminus. Here we find that the proposed C-terminal D-box of AURKA does not act as a degron and instead mediates essential structural features of the protein. In living cells, as previously reported *in vitro*, the N-terminal intrinsically disordered region (IDR) of AURKA containing the A-box is sufficient to confer FZR1-dependent mitotic degradation. Both *in silico* and *in cellulo* assays predict the QRVL Short Linear Interacting Motif (SLiM) of the A-box to be a phospho-regulated D-box. We propose that degradation of full-length AURKA additionally depends on an intact C-terminal domain because of critical conformational parameters permissive for both activity and mitotic degradation of AURKA.

**Summary blurb:** AURKA degron motifs are redefined to show that the so-called N-terminal ‘A-box’ is in fact a D-box, and the so-called ‘D-box’ in the C-terminus is not a degron but a motif critical for the active, degradable conformation of AURKA

## Introduction

The control of mitotic exit is a paradigm for cellular regulation through targeted proteolysis. The process is orchestrated by a multi-subunit ubiquitin ligase (E3) complex, the Anaphase-Promoting Complex (APC/C), which utilizes two WD40 domain co-activator paralogues contributing to substrate recognition, CDC20 and Cdh1/FZR1 (henceforth referred to as FZR1). E3s recognise their targets through substrate motifs called degrons, and although some degrons conform to a robust consensus (for example the di-phospho-degrons recognized by SCFβTRCP), most are rather ill-defined and can usefully be defined as Short Linear Interacting Motifs (SLiMs) present in intriniscally disordered regions of proteins, or IDRs. SLiMs are proposed to adopt transient structures to mediate weak protein-protein interactions such as those between substrate and E3 ligase (Van Roey et al. 2014). Such structures have now been solved by cryoEM to identify docking sites on the APC/C for degrons known as the D-box (Destruction-Box), KEN motif and ABBA motif (Chao et al. 2012; He et al. 2013; Chang et al. 2015; Brown et al. 2016). The first D-box was identified in the N-terminal IDR of cyclin B1 (Glotzer, Murray, and Kirschner 1991), subsequently a large number have been found in other APC/C substrates, most fitting the general consensus RxxLxxxxx, although a number of variants have been identified that lack either R at position 1 (P1) or L at P4 (He et al. 2013; Davey and Morgan 2016). The KEN motif is extremely common in the proteome, first identified in CDC20 (Pfleger and Kirschner 2000) when it was thought to be specific to substrates targeted by the FZR1-activated version of the APC/C but now revealed to be a universal APC/C degron that docks onto the upper surface of the WD40 propellor of either CDC20 or FZR1 (Chao et al. 2012; He et al. 2013). The ABBA motif has been identified in BubR1 and cyclin A and confers increased affinity that is essential for control of the mitotic checkpoint (Di Fiore et al. 2015; Qin et al. 2016).

It seems likely that a combination of degrons for multivalent docking at the KEN receptor and D-box receptor (DBR) in the APC/C are required to generate selectivity in substrate targeting amongst the large number of degrons motifs present in the proteome, and to generate the increased affinity of APC/C-substrate interaction required for efficient ubiquination (Lu, Wang, and Kirschner 2015). As discussed in (Davey and Morgan 2016), lack of strict conservation of degron sequences can be explained by multivalency of degron-E3 interactions and participation of residues outside the consensus. These features go hand in hand with flexibility of IDRs, and flexibility in sequences surrounding the degron are essential for substrate lysines to be able to mount nucleophilic attack on a nearby ubiquitin thioester linkage. The lack of sequence conservation between degrons has contributed to historic confusion in the field, with some atypical degrons described as ‘novel’ being subsequently redefined as variants on known degrons (i.e. they dock to the known receptor sites on the APC/C) (He et al. 2013; Davey and Morgan 2016). Indeed a systematic study of sixteen APC/C^FZR1^ substrates in budding yeast concluded that all of them depended on the DBR of FZR1 for their degradation, even when they did not have an obvious D-box (Qin et al. 2016). One candidate atypical D-box that has not been further investigated is the so-called ‘A-box’ of AURKA (Littlepage and Ruderman 2002).

AURKA is a substrate of the APC/C during anaphase that, unlike other anaphase substrates, is specific to the FZR1-bound form of APC/C (Honda et al. 2000; Castro, Arlot─ Bonnemains, et al. 2002; Kitajima et al. 2007; Floyd, Pines, and Lindon 2008; Min et al. 2015). Potential degrons in AURKA have been defined mostly through *in vitro* studies, with deletion or mutation of either the ‘A-box’ motif or the putative D-box shown to stabilize AURKA against mitotic degradation in one or more assays *in vitro* or in living cells (summarized in Table 1). There is widespread acceptance in the literature that destruction of full-length AURKA depends upon an N-terminal A-box and C-terminal D-box.

**Table 1.** Summary of AURKA degrons described in the scientific literature

| reported AURKA degron |  |  | Evidence *in vitro* | Evidence in cells | Notes |
| --- | --- | --- | --- | --- | --- |
| Name | position |  |  |  |  |
|  | Xenopus | Human |  |  |  |
| D-box<sup>1</sup> | R378 |  | Arlot-Bonnemains 2001 |  | RxxL>RxxI stabilizes in CSF extract assay |
|  |  |  | Castro 2002a |  | RxxL>AxxA stabilizes and reduces FZR1-dependent ubiquitination |
|  |  |  | Littlepage 2002a |  | 1-386 or RxxL>AxxL are stable versions |
|  |  | R371 | Crane 2004 |  | *in vitro* degradation assays |
|  |  |  |  | Kitajima 2007 | sensitivity to FZR1 overexpression |
| KEN<sup>2</sup> | K6 |  | Arlot-Bonnemains 2001<br>Littlepage 2002a |  | very small effect on degradation, not thought to be a degron |
|  |  | K5 | Crane 2004 |  | very small effect on degradation, not thought to be a degron |
|  |  |  |  | Min 2014 | identified as ubiquitinated residue |
| A-box | 34-69 |  | Littlepage 2002a |  | 1-136 sufficient for degradation |
|  | 44-55 |  | Castro 2002b |  | A-box behaves as 'D-box Activating Domain' (DAD) |
|  |  | 32-65 | Crane 2004 |  | *in vitro* degradation assays |
|  |  |  |  | Floyd 2008 | live cell mitotic degradation |
| QRVL<sup>2</sup> |  | Q45 | Crane 2004 |  | *in vitro* degradation assays |
|  |  |  |  | Kitajima 2007 | sensitivity to FZR1 overexpression. |
| Phospho-site<sup>3</sup> | S53 |  | Littlepage 2002a<br>Littlepage 2002b<br>Castro 2002b |  | phospho-mimic versions stable in *in vitro* degradation assays. |
|  |  | S51 | Crane 2004 |  | phospho-mimic versions stable in *in vitro* degradation assays. |
|  |  |  |  | Kitajima 2007<br>Lindon 2016 | sensitivity to FZR1 overexpression.<br>live cell mitotic degradation. |

1 Also shown for AURKB D-box *in vitro* in (Stewart and Fang 2005) but not in (Nguyen et al. 2005).
2 Identified as degron in AURKB (Nguyen et al. 2005).
3 pS51 found in proteomic studies as *bone fide* phosphorylation site (Daub et al. 2008; Dulla et al. 2010; Plotnikova et al. 2010).

However it is also well known from the crystal structure of the AURKA kinase domain that the putative D-box is located in a highly structured domain - inconsistent with the definition of a SLiM – and that its conserved RxxL residues are structurally buried (Bayliss et al. 2003). The proposed degron would therefore be inaccessible to APC/C binding unless ubiquitination of AURKA were preceded by an unfolding step, rendering problematic the designation of this motif as a degron, despite the *in vitro* evidence of degron function.

Here we describe a study undertaken to provide a more complete characterization of AURKA degrons. We report that the proposed C-terminal D-box does not display properties of a degron and instead mediates structural features of the protein essential for its normal function. We show that in living cells, as has previously been reported *in vitro* (Littlepage and Ruderman 2002), the N-terminal IDR of AURKA is sufficient to confer FZR1-dependent mitotic degradation, and that both *in silico* docking experiments and live cell degradation assays predict the so-called A-box to be an N-terminal D-box. Nonetheless degradation of full-length AURKA additionally depends on an intact C-terminal domain, because of critical conformational parameters. We propose an explanation of how mitotic degradation of AURKA is sensitive to the conformational dynamics of the substrate.

## Results & Discussion

A number of publications identify putative degrons in AURKA through studies on both Xenopus and human proteins (Table 1). Since the existence of a D-box within the structured kinase domain of AURKA has been called into question (Davey and Morgan 2016; Lindon, Grant, and Min 2015), we decided to look more closely at the putative D-box status of this motif through a combined analysis of structure, function and degradation of different AURKA mutants.

The structure of hsAURKA kinase domain (122-403) was examined in PyMol and variations in free energy resulting from different point mutations in the putative D-box (R_371_xxL) or in an adjacent D-box-like motif (R_375_xxL) were calculated using FoldX3 software (Figure 1). The molecular structure of the putative D-box shows L374 fitted into the hydrophobic aliphatic pocket on the kinase domain and a salt bridge established between R371 and conserved residue E299 (Figure 1A). Gibbs free energy variations (ΔΔG) for the protein folding state predict that the RxxL>AxxA substitution frequently used to test for D-box function is strongly destabilizing to the structure (R371A/L374A, ΔΔG = 5.8 kcal mol^- 1^). R371A contributes most of the free energy variation (4.9 kcal mol^-1^) with L374A less destabilizing (ΔΔG = 2.6 kcal mol^-1^). The conserved substitution L374I is predicted to destabilize least of all (ΔΔG = 1.6 kcal mol^-1^) (Figure 1B).

**Figure 1:**
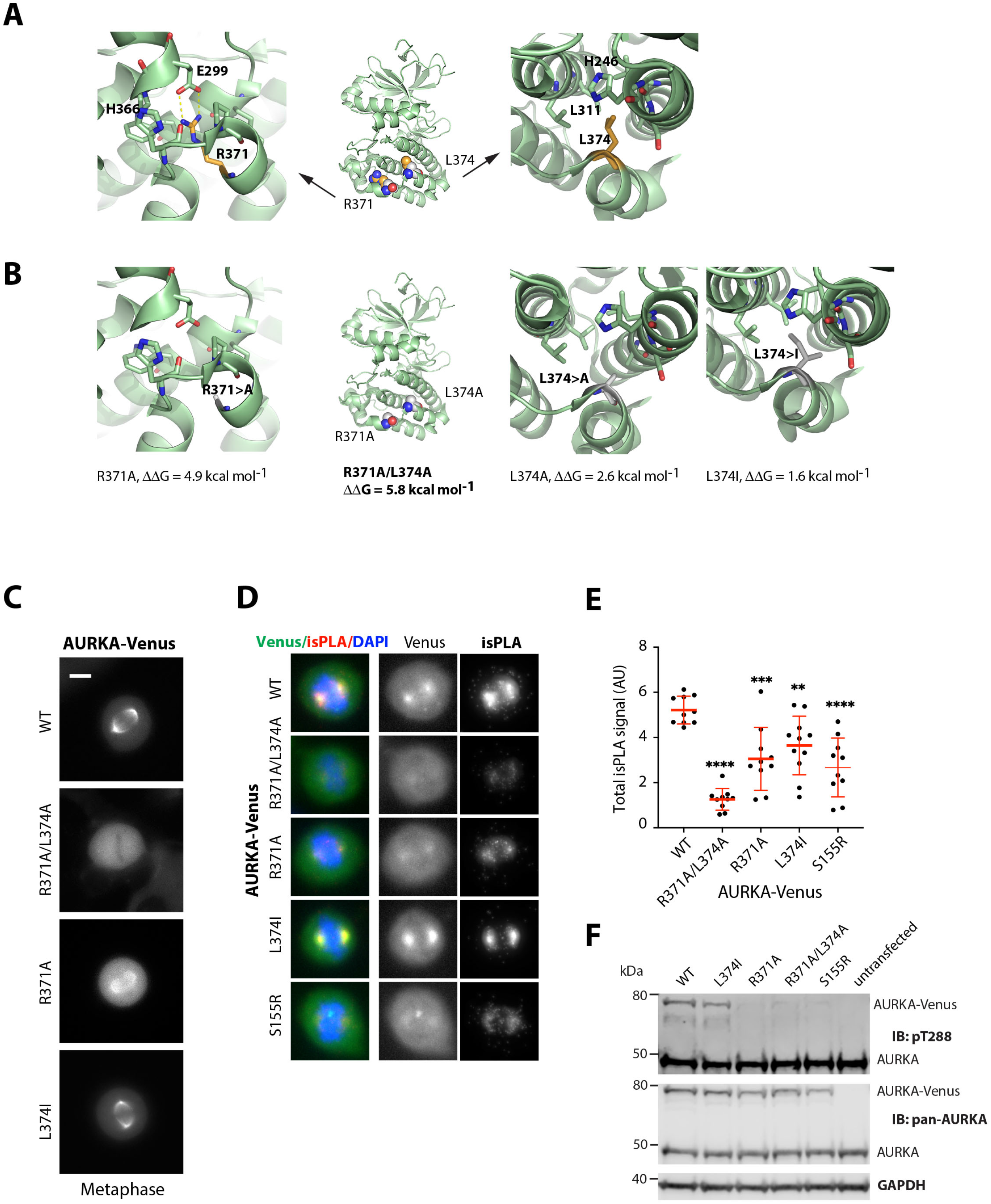
The C-terminal R_371_xxL motif of AURKA plays a critical role in folding and function. **A-B** *In silico* testing of the C-terminal D-box-like motif. **A** R_371_xxL motif is buried within the kinase domain. Arginine residue R371 (orange) establishes salt bridges with conserved glutamic acid residue E299 (green). Leucine L374 fits into the hydrophobic aliphatic pocket on the kinase domain. **B** interactions shown in **A** are lost in structures predicted for R371A and L374A substitutions. Gibbs free energy variations (ΔΔG = ΔGmut-ΔGwt) for the protein folding state were predicted using FoldX3 software and show that R371A and L374A substitutions are more strongly destabilizing to the structure than the conserved substitution L374I. **C-E** Characterisation of different versions of Venus-tagged AURKA in human U2OS cells. **C** Panels showing localization of AURKA-Venus in live cells during interphase and at mitosis. **D-E** Interaction with TPX2 assayed by isPLA (see also Supplementary Figure 2). **D** isPLA signal reveals AURKA-Venus-TPX2 interaction on mitotic spindles. **E** Total isPLA signal was quantified per mitotic cell and corrected for background. Data for each condition (n ≥ 10) was plotted in a scatter plot with bar and whiskers to indicate mean and SDs. Normality was verified by D’agostino and Pearson test and each mutant compared to the wild type by Ordinary One-way Anova. **, p ≤ 0.01; ***, p ≤ 0.001; ****, p ≤ 0.0001. Results representative of two identical repeats of the experiment. **F** Different versions of AURKA-Venus from transiently transfected U2OS cells arrested in mitosis were probed for pT288-AURKA by immunoblot. Blot representative of two experiments.

We examined the localization of YFP- or Venus-tagged versions of AURKA in living cells and found that versions with structurally destabilizing substitutions in the R_371_xxL motif did not behave like the wild-type (WT) protein, being excluded from (R371A/L374A) or only weakly present (R371A) on the microtubules of the mitotic spindle (Figure 1C). By contrast the less destabilizing substitution (RxxI) did not affect localization in mitotic cells. The ‘non-degradable’ version of AURKA (ΔA-box) generated by deletion of the N-terminal A-box degron (Littlepage and Ruderman 2002; Castro, Vigneron, et al. 2002) has previously been shown to localize as the WT protein (Floyd, Pines, and Lindon 2008; Lindon, Grant, and Min 2015). Since some versions were expressed at lower levels than the WT protein, we simultaneously depleted endogenous protein to eliminate competition from unlabelled endogenous AURKA to confirm that R371A/L374A does not localize to the spindle even when in excess over the endogenous protein (Supplementary Figure S1).

The various subcellular localizations of the Aurora kinases are known to depend on their binding partners, with the best characterized partner of AURKA, TPX2, responsible for its localization to the mitotic spindle (Kufer et al. 2002). We therefore examined the interaction of different versions of AURKA-YFP with endogenous TPX2, including the known loss of interaction mutant S155R (Bibby et al. 2009) using an in situ Proximity Ligation Assay (isPLA). As expected we found that the R371A/L374A version of AURKA, which does not localize to the mitotic spindle, does not interact with TPX2 (Figure 1D, Supplementary Figure S2). Quantification of isPLA signals confirmed that the amount of TPX2 interaction measured for different R_371_xxL mutants (Figure 1E) correlated with the degree of spindle localization observed (Figure 1D). Probing the same versions of AURKA with antibody against the phospho-T288 epitope by immunoblot confirmed that detection of auto-activated kinase is severely compromised in R371A/L374A, as predicted for the S155R version (Figure 1F). We concluded that perturbed folding of the C-terminal region of its kinase domain prevents AURKA folding and function required for interaction with TPX2.

Next, we compared our panel of putative D-box substitutions for their effect on mitotic degradation of AURKA-Venus using a fluorescence timelapse assay. The R371A/L374A version was resistant to mitotic degradation, as is the ΔA-box version lacking the previously characterized N-terminal degron sequence located between resides 32 and 66 (Figure 2A). The R371A substitution, with partial destabilizing effect on AURKA, showed partial resistance to mitotic degradation. L374I, with the lowest ΔΔG, had no effect on mitotic degradation of the protein (Figure 2B). Plotting the amount of degradation (as percentage of each version of AURKA remaining 60 minutes after anaphase) against the calculated ΔΔG of folding revealed a significant inverse correlation of the two values (Figure 2C). These results suggested that perturbed folding could be responsible for both the deficiency in interaction with TPX2 *and* deficiency in mitotic degradation of AURKA R_371_xxL (so-called ‘D-box’) mutants, and did not allow us to conclude whether or not the R_371_xxL motif was a *bona fide* degron.

**Figure 2:**
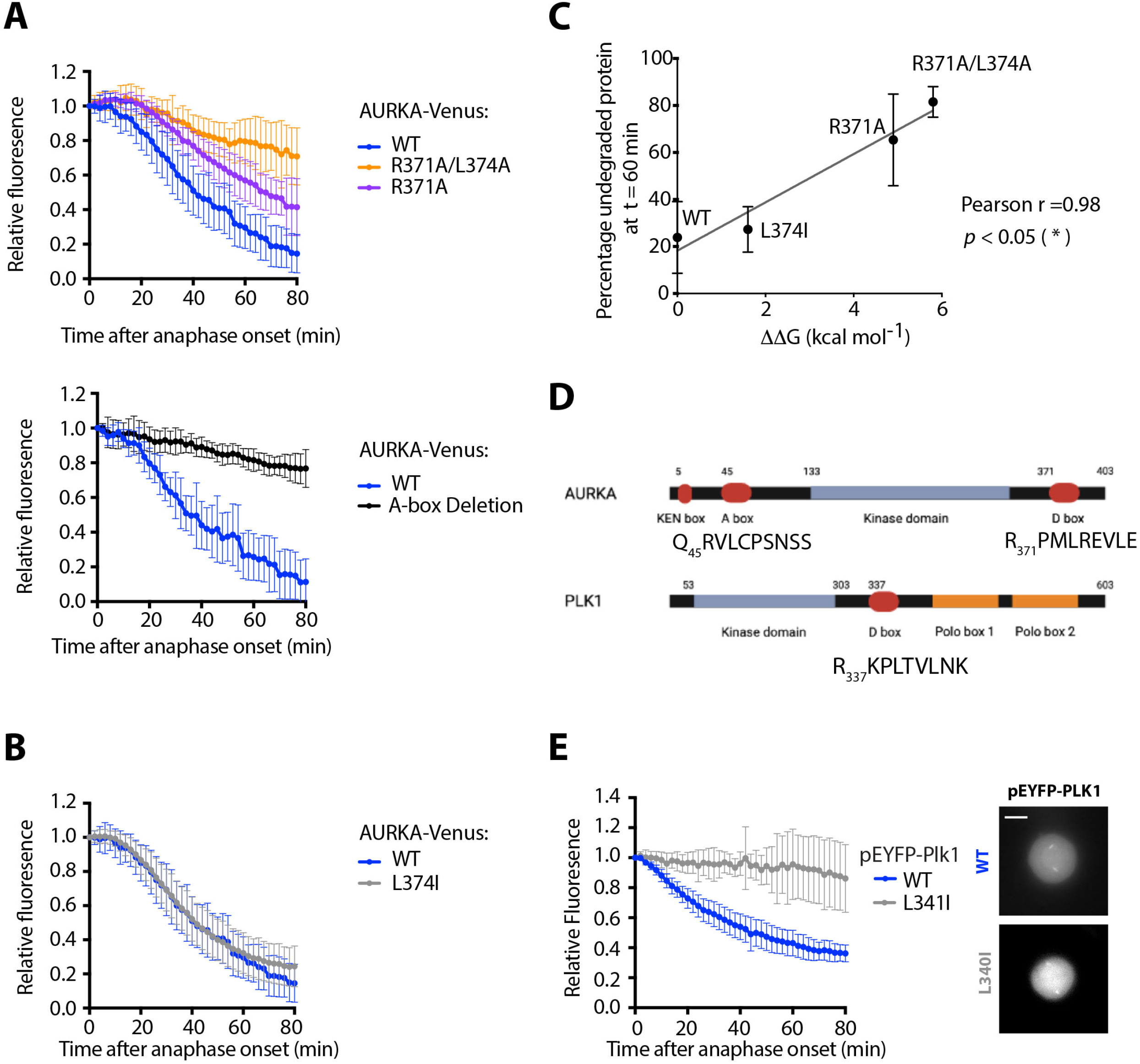
The R_371_xxL motif is not a D-box. **A-B** *In cellulo* mitotic degradation assays of AURKA-Venus. Graphs show quantified Venus levels from fluorescence timelapse imaging of single cells undergoing mitotic exit. Venus levels from individual cells are normalized against the Venus level at anaphase onset and *in silico* synchronized so that mean & SDs can be plotted for each version of AURKA-Venus. **A** Mutations predicted to cause disruption of the C-terminus block AURKA degradation during mitotic exit: R371A/L374A (top graph, n ≥ 10 cells, pooled from two experiments) blocks degradation of AURKA-Venus in a similar way to deletion of the N-terminal A-box (bottom graph, n ≥ 6 cells, from a single experiment)(Floyd, Pines, and Lindon 2008). **B** The conservative substitution L374I has no effect on kinetics of degradation of AURKA-Venus (n ≥ 10 cells, pooled from two experiments). **C** Correlation plot for percentage degradation of each version of AURKA-Venus versus predicted ΔΔG of substituted residues in R_371_xxL. **D** Schematic of known and proposed degrons in AURKA and Plk1. **E** *In cellulo* mitotic degradation assays of YFP-Plk1 WT and L341I version (n ≥ 10 cells, pooled from two experiments). The L>I substitution at P4 of the R_337_xxL motif abrogates degradation at mitotic exit whilst having no effect on localization of the protein (right hand panels).

Therefore we investigated further whether the different R_371_xxL mutations showed a pattern of mitotic degradation consistent with the designation of this motif as a D-box. Our observations that Ala substitutions at P1 and P4 of the putative D-box stabilized AURKA against mitotic degradation were consistent with the known literature on D-box motifs. However, our finding that L>I at P4 (L374I) had no effect on mitotic degradation was not consistent. An isoleucine sidechain at P4 position of a D-box would be predicted to disrupt binding to the DBR due to the shifted methyl in isoleucine that reduces the Van der Waals contribution of this residue to the interaction in the hydrophobic pocket, and should be sufficient to stabilize a substrate against APC/C-mediated degradation. Indeed, this had previously been reported for mutation of the equivalent P4 residue in Xenopus Aurora A (L381I) in *in vitro* degradation assays (see Table 1). We tested the prediction on a well-known anaphase substrate of APC/C, Polo-like kinase 1 (Plk1), whose degradation is critically dependent on a single D-box (R_337_xxL, Figure 2D) (Lindon and Pines 2004). We introduced the single L>I substitution at P4 of the Plk1 D-box and found that this version, L340I, was unable to mediate any mitotic degradation of correctly localized Plk1 (Figure 2E). Taken together, our findings support the prediction that L>I substitution at P4 abrogates D-box function and that – since in our experiments AURKA L374I is degraded normally – the R_371_xxL motif in human AURKA is unlikely to function as a D-box degron.

Since the C-terminal RxxL motif of AURKA was probably not a D-box, we investigated whether the N-terminal IDR would be sufficient – as well as necessary - for mitotic degradation in live cell assays. The A-box (32-66) has been shown in a number of studies to be essential for AURKA degradation in a manner that depends critically on the strongly conserved Q_45_RVL SLiM and on the phosphorylation status of S51 (Table 1). The N-terminal K_5_EN motif contributes to degradation, although since K5 is a ubiquitination site during mitotic exit (Min, Mayor, and Lindon 2013), it is not clear whether this motif acts as a degron, provides the ubiquitin receptor in the substrate, or both. We first tested AURKA(1-133) fused to GFP in live cell assays (Supplementary Figure S3) and subsequently AURKA(1-67) fused to mNeonGreen (mNeon), after confirming that the mNeon-tagged version was degraded more efficiently than the GFP-tagged version (as reported in Khmelinskii et al. 2016). We found that AURKA(1-67) was sufficient to direct anaphase-specific, FZR1-dependent degradation of fluorescent protein tags (Figure 3A). The full-length protein was degraded more efficiently than the N-terminus alone, but we could conclude nonetheless that 1-67 of AURKA contains degrons required for its targeted destruction at mitotic exit. We further tested, in the context of AURKA(1-67), features contributing to specificity of AURKA targeting at mitotic exit, finding that both K_5_EN and Q_45_RVL SLiMs were required for degradation (Min, Mayor, and Lindon 2013)(Figure 3B). Therefore AURKA (1-67) recapitulates the known regulation of degradation of the full-length protein.

**Figure 3:**
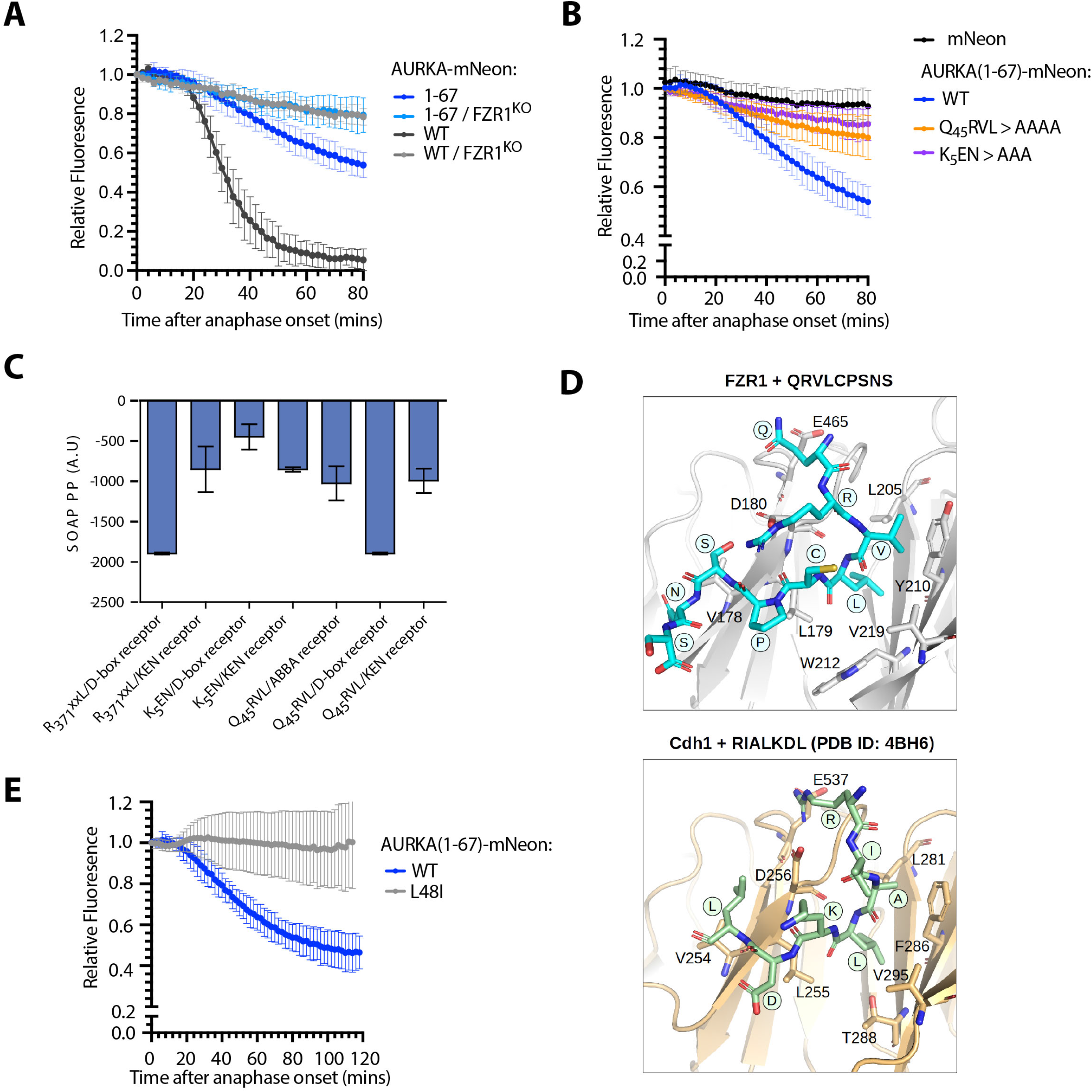
The Q_45_RVL motif within the AURKA A-box displays properties of a D-box degron. **A-B** *In cellulo* mitotic degradation assays for full-length AURKA-mNeon and AURKA(1-67)-mNeon expressed in U2OS or FZR1^KO^ U2OS cells. mNeon levels from individual cells are normalized against the anaphase onset level and *in silico* synchronized so that mean and SDs can be plotted for each protein (n ≥ 20 cells pooled from ≥ 3 experiments). Degradation curves show that (**A**) residues 1-67 are sufficient for mitotic exit-specific FZR1-dependent degradation and (**B**) mitotic degradation of AURKA(1-67)-mNeon depends on SLiMs at K_5_ and Q_45_RVL. **C** Energetics of *in silico* docking of proposed AURKA degrons into known binding pockets on FZR1, scored by SOAP-PP, using FlexPepDock server. **D** A-box (QRVLCPSNS) peptide docked to the *H*.*s*. FZR1 DBR (top panel), modelled upon structure PDB 4BH6 that shows *S*.*c*. Cdh1 bound to D-box peptide (RIALKD). PDB 4BH6 is shown for comparison in the bottom panel (He et al. 2013). **E** *In cellulo* mitotic degradation assays of AURKA-Venus WT and L48I (L>I at P4 of A-box motif, n = 26 cells pooled from two experiments).

We turned to an *in silico* docking approach to examine whether the atypical Q_45_RVL degron might bind one of the known degron receptor sites on FZR1. We docked the peptide Q_45_RVLCPSNS into the sites on FZR1 (D-box, KEN, ABBA) identified from the cryo-EM structure of FZR1 WD40 domain bound to the pseudosubstrate domain of Acm1 (PDB:4BH6, He et al. 2013)(Figure 3C, Supplementary Figure S4), using the FlexPepDock server (Raveh, London, and Schueler-Furman 2010). The optimized pose at each site was compared with binding of a cognate degron and the Statistically Optimized Atomic Potential (SOAP) assigned to each interaction. This revealed that the Q_45_RVL peptide favoured interaction at the D-box pocket compared to other sites, and that docking of Q_45_RVL at this site was energetically comparable to docking of D-box peptides (Figure 3C). Comparison of the A-box with a panel of D-boxes showed that it docks with an affinity within the range of known D-boxes, although with reduced affinity compared to more canonical D-boxes (Supplementary Figure S4). We note that *in silico* docking to FZR1 alone ignores any contacts made between substrate and APC10 that probably contribute to the affinity of substrate for the D-box binding pocket (Qin et al. 2019) and therefore we may have underestimated the likely preference of Q_45_RVL for the DBR.

The Q_45_RVLCPSNS peptide can be docked in a similar pose to canonical D-box peptides, such that the peptide adopts a 90º turn at P4 with the leucine sidechain extending into the hydrophobic cleft in FZR1 identified by He et al. 2013 (Figure 3D), and consistent with L at P4 being the most critical residue of the D-box (He et al. 2013; Davey and Morgan 2016). We tested the importance of the leucine sidechain by making the L>I substitution at this position, the ‘D-box test’ predicted to disrupt docking at the DBR (and shown to block degradation of Plk1 in Figure 2D). The L48I version of AURKA(1-67) was resistant to anaphase degradation, consistent with the hypothesis that the so-called A-box (Q_45_RVL) is a D-box (Figure 3E).

Given the conservation of R at P1 in most D-boxes, how does the AURKA Q_45_RVL dock efficiently at the DBR? *In silico* docking predicts that some of the electrostatic interactions with E465 and D180 of FZR1, usually made by R at P1 of canonical D-boxes, could be replaced by R at P2 (Figure 4A). In He et al. (2013) it is argued that contacts made by P7 of the D-box could compensate for lack of R at P1. In AURKA, P7 of the Q_45_xxL D-box is occupied by S51, which contacts D180 in the docking pose shown in Figure 4A. We note that in alternative poses, where the P7 residue does not contact D180 we observe steric clashes between D-box residues at P8 and P9, and APC10. A block on degradation caused by phosphomimetic substitution at S51 has been a consistently observed feature of AURKA mitotic degradation over two decades (Table 1, Figure 4B). We confirmed in cell-based assays that S51D abrogates ubiquitination of full-length AURKA-Venus as efficiently as mutation of Q_45_RVL or deletion of the entire A-box (32-66), pointing to a critical role of the residue in P7 in docking at the DBR (Figure 4C). We then compared *in silico* docking of wild-type A-box peptide (S at P7) with its ‘non-degradable’ version (D at P7) version, computing free energy values using FoldX3. This analysis confirmed increased binding energy of the mutant (Figure 4D), resulting from electrostatic repulsion between the negative charge at P7 and D180 of FZR1 (Figure 4A). The ∼2.0 kcal/mol difference that we observed translates into a predicted ∼30 fold increase in K_d_ and is consistent with the inhibition of mitotic exit degradation by S51D substitution. Our *in silico* docking model therefore explains known features of AURKA degradation and assigns the ‘A-box’ degron to the category of phospho-regulated D-boxes alongside securin, CDC6, KIFC1 (Holt 2012; Singh et al. 2014). We note that a recent study from the Barford lab comparing cryo-EM structures of D-boxes in Cyclin A reveals that canonical D-box D1 (RxxL) binds in a different mode to a newly identified non-canonical D-box D2 (VxxL) and that, in *in vitro* ubiquitination assays, D2 behaves as a stronger D-box degron than D1 (PDB:6Q6H, S. Zhang, Tischer, and Barford 2019). Overall, our findings are consistent with the idea that specific degron properties can be encoded in non-canonical D-boxes and that Q_45_RVL is a APC/C^FZR1^-specific D-box.

**Figure 4:**
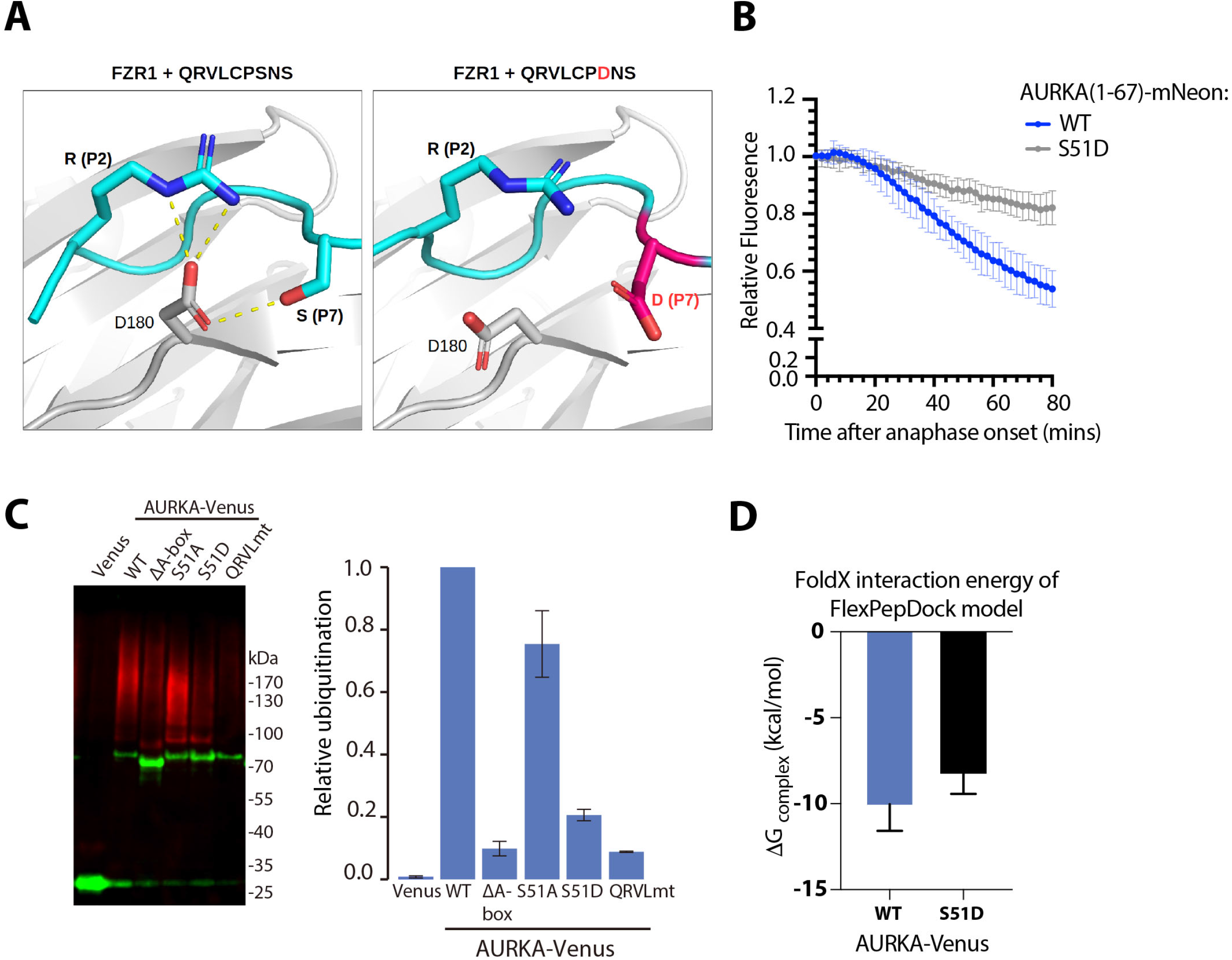
Modelling of the Q_45_RVL motif at the DBR explains the role of AURKA S51 phosphorylation. **A** Docking of Q_45_RVLPSNSS peptide on FZR1. Predicted pose for QRVL in the DBR by *in silico* docking, showing orientation of the P4 Leucine and novel contacts afforded at P2 and P7. **B** *In cellulo* mitotic degradation assays of AURKA-Venus WT and S51D. **C** Ubiquitination of wild-type and different versions of AURKA carrying mutations in the A-box, transiently expressed Venus-tagged proteins were purified from U2OS cells synchronized in mitotic exit and blotted for GFP (in green) and ubiquitin conjugates (FK1 antibody, in red). Relative ubiquitination was plotted as the ratio of ubiquitin-conjugated:unmodified protein, normalized against the wild-type protein, error bars show SDs from 3 repeats of the experiment. **D** Free energy values for WT and S51D peptides computed using FoldX3. *In silico* docking models were rebuilt using mutant peptide QRVLCPDNS, models were scored with FoldX3 and the average binding free energies of 10 models for each were plotted. The higher binding energy of the mutant is significant according to Mann-Whitney test (p = 0.0147).

If all sequences required for APC/C^FZR1^-mediated degradation of AURKA reside in its N-terminal IDR, then why does mutation in the C-terminus stabilize the full-length protein? In this study we have shown (Figure1) that the R_371_xxL motif is required for interaction of AURKA with TPX2. Is TPX2-mediated activation of AURKA therefore required for it to be a target of APC/C? We tested this question by examining mitotic degradation of AURKA S155R-Venus (which interacts only weakly with TPX2, Figure 1E). We found that S155R was degraded with identical kinetics to the wild-type protein (Figure 5A) and concluded that interaction with TPX2 *per se* is not required for AURKA-Venus degradation. Rather, we propose that interaction of AURKA with TPX2 and with APC/C-FZR1 both depend upon a conformational state blocked by disruption of the R_371_xxL motif in the C-terminal helix of the kinase domain.

**Figure 5:**
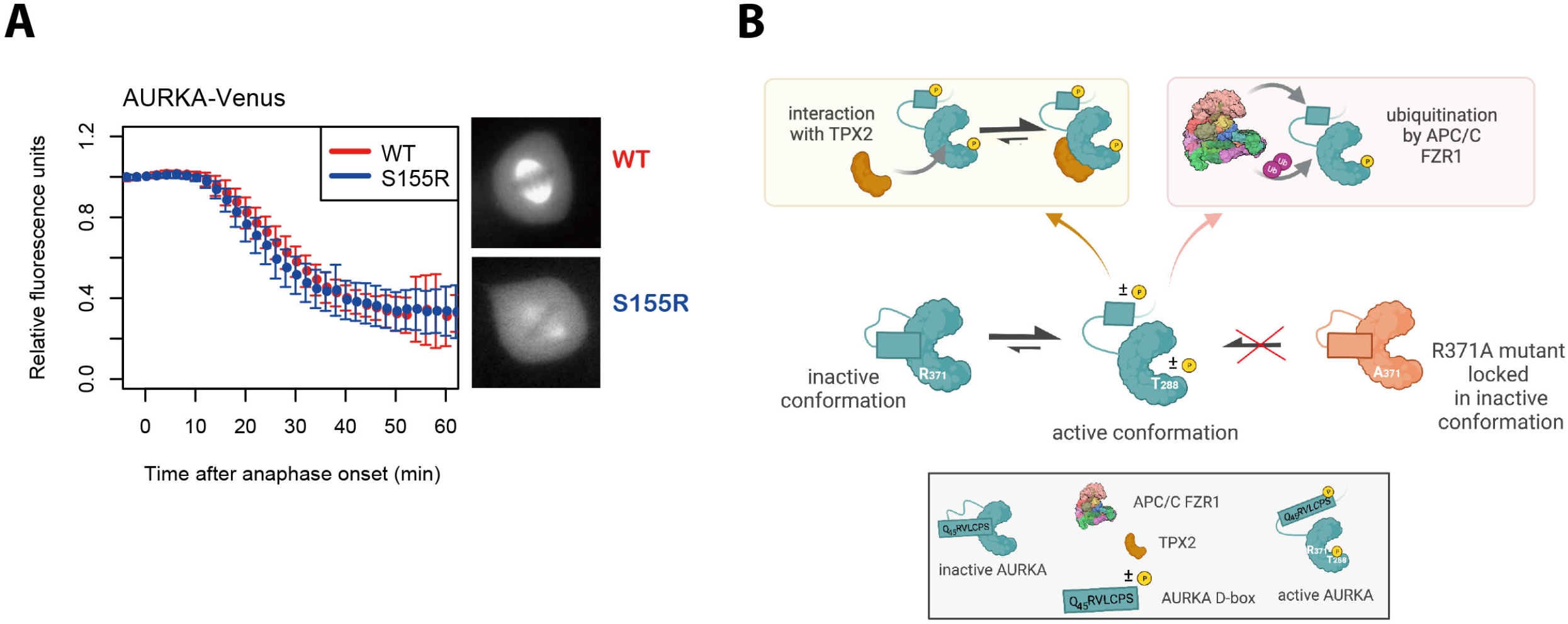
R_371_xxL motif plays a critical role in conformational regulation of AURKA. **A** *In cellulo* mitotic degradation assays of AURKA-Venus WT and S155R. Venus levels from individual cells are normalized against the anaphase onset level and *in silico* synchronized so that mean and SDs can be plotted for each protein (n ≥ 10 cells, representative of two experiments).**B** Schematic proposing that the link between R371 and degradability of AURKA are mediated by a conformational step that simultaneously activates AURKA (leading to phosphorylation on T288) and makes it degradable by APC/C. The Q_45_RVL motif is ‘buried’ in the autoinhibited state of the WT protein (green) and once released is autoregulated by phosphorylation on S51. The R371A mutant (orange) is unable to undergo the critical conformational step required for both activation and degradation.

Taken together, our results indicate that mutation in R_371_ prevents the active, degradable conformation of AURKA. An auto-inhibitory state involving interaction between the kinase domain and the N-terminus of AURKA has previously been described (Y. Zhang et al. 2007; Bai et al. 2014) and studies with a FRET-based conformational sensor confirm that the relative configuration of N- and C-terminus is altered upon activation of the kinase (Bertolin et al. 2016). Since interaction with APC/C-FZR1 occurs through an N-terminal degron motif, we propose that the Q_45_xxL D-box is ‘buried’ in the inactive conformation, which may be the previously described auto-inhibited state mediated through interaction between N- and C-terminal domains (Figure 5B). Intriguingly, a heterotetrameric structure for AURKA-TPX2 captured by the Kern lab (Zorba et al. 2014) shows AURKA as a dimer stabilized by a ‘dimer swap’ interaction of R371 residue with E299 in a second AURKA molecule. That is, the usual stabilizing interaction between R371 and E299 can occur in intermolecular fashion instead of intramolecular and the authors propose dimerization as a necessary step in auto-activation of the kinase (promoting autophosphorylation, allowing release of the autoinhibited state and interaction with TPX2). The most recent published studies of AURKA have described redox regulation of AURKA activity through oxidation or inhibitory CoA-lation of a cysteine residue in the kinase activation loop (Byrne et al. 2020; Tsuchiya et al. 2020) also implicated in AURKA dimerization in vitro (Zorba et al. 2014; Tsuchiya et al. 2020), indicating that there is still much to learn of the conformational complexities of AURKA.

Our own findings are consistent with the idea that R371 facilitates an activation step of AURKA and reconciles a large amount of disparate and sometimes contradictory literature on the role of the C-terminal R_371_xxL (historically referred to as the ‘D-box’) in the activity, function and degradation of AURKA. In summary, we present evidence that the previously assigned ‘D-box’ (R_371_xxL) of AURKA is not a degron, and that the previously named ‘A-box’ degron (Q_45_RVL) is the *bona fide* AURKA D-box. Given the timing and dependencies of AURKA degradation, we propose that the A-box should be considered an APC/C-FZR1-specific D-box.

## Materials and Methods

### In silico mutagenesis and docking

Variation in free energy of folding upon point mutations was estimated using the FoldX programme (Guerois, Nielsen, and Serrano 2002). Docking was performed using FlexPepDock to return a docked model for each peptide tested. Where indicated, these were scored for comparison using the Statistically Optimized Atomic Potential for protein-protein docking (SOAP-PP) assigned to each interaction (Dong et al. 2013). Docking simulations at the DBR were filtered for L4 in same position as for Acm1 bound to S. cerevisiae Cdh1 (PDB code 4BH6). No positional restraints were applied when docking at the KEN/ABBA sites.

### Plasmids

pVenus-AURKA-N1 has been described in previous publications (Min, Mayor, and Lindon 2013). N-terminal fragments 1-133 and 1-67 of AURKA were cloned into pEGFP-N1, with EGFP substituted by mNeon-HA for later experiments. Point mutagenesis on AURKA was carried out using standard techniques, all cloning details are available upon request.

### Cell culture, synchronization and transfection

U2OS cells were cultured in DMEM supplemented with 10% FBS, 200 µM Glutamax-1, 100 U/ml penicillin, 100 µg/ml streptomycin, and 250 ng/ml fungizone (all from ThermoFisher Scientific) at 37ºC in humidified atmosphere containing 5% CO_2_. Cells were transfected using a Neon electroporator (ThermoFisher Scientific) and seeded onto 8-well microscope slides (Ibidi) for live cell imaging or onto coverslips for PLA assays.

For synchronization in mitosis, cells were treated for 12 h with 10 μM STLC (Tocris Bioscience) to trigger the Spindle Assembly Checkpoint (SAC). For mitotic exit synchronizations cells synchronized with STLC were collected by shake-off and released by 70 minute treatment with 10 μM ZM447439 (Generon, UK), an inhibitor of AURKB which ablates SAC arrest.

### Fluorescence Microscopy of living cells

A few hours before live cell imaging of Ibidi slides, the culture medium was changed to L15 supplemented with 10% FBS, antibiotics and antimycotics. Images were acquired using an automated epifluorescence imaging platform composed of Olympus IX83 motorized inverted microscope, Spectra-X multi-channel LED widefield illuminator (Lumencor, Beaverton, OR, USA), Optospin filter wheel (Cairn Research, Faversham, UK), CoolSnap MYO CCD camera (Photometrics, Tuscon, AZ, USA), automated XY stage (ASI, Eugene, OR, USA) and climate chamber (Digital Pixel, Brighton, UK) and controlled using Micro-Manager (Edelstein et al. 2014). For mitotic degradation assays, multiple fields containing Venus-, EGFP- or mNeon-positive prophase and metaphase cells were selected. Images were then acquired every 2 mins for 2 hours using appropriate filter sets. Fluorescence levels were measured using ImageJ and degradation curves plotted in GraphPad Prism (San Diego, CA, USA), *in silico* synchronized to anaphase onset for each construct tested.

### In situ Proximity Ligation Assay (isPLA)

U2OS cells were transfected with AURKA-Venus wild-type or mutant and seeded on coverslips. After 24 hours cells were fixed with 4% PFA and processed for isPLA using Duolink® In Situ Detection Orange (Sigma-Aldrich) according to manufacturer’s instructions, using primary antibodies against GFP (Mouse mAb clone, #11814460001, Roche) and TPX2 (rabbit polyclonal antibody, Novus Biological). Epifluorescent images were acquired on the widefield imaging platform described above, as stacks of 500 nm step with 2×2 bin, using appropriate filter sets and 40X NA 1.3 oil objective, and exported as maximal intensity projections in ImageJ (http://rsb.info.nih.gov/ij/; National Institutes of Health, Bethesda, MD). Integrated intensities were measured and total isPLA signal quantified per mitotic cell and corrected for background.

### Immunoblotting

Cells were lysed in in 1% Triton X─ 100, 150 mM NaCl, 10 mM Tris–HCl at pH 7.5 and EDTA-free protease inhibitor cocktail (Roche), and PhosSTOP™ inhibitor for phosphatase (Sigma-Aldrich). After 30 min on ice, the lysate was centrifuged at 14,000 rpm (4°C) for 10 min. For immunoblotting, an equal amount of protein (20 μg) were loaded into SDS-PAGE 4-12% pre-cast gradient gels. Proteins were transferred to Immobilon-P or Immobilon-FL membranes using the XCell IITM Blot Module according to the manufacturer’s instructions. Membranes were blocked in PBS, 0.1% Tween-20, 5% BSA and processed for immunoblotting. Primary antibodies for immunoblot were as follows: AURKA (1:1000; mouse mAb clone 4/IAK1, BD Transduction Laboratories), phospho-Aurora A (Thr288)/Aurora B (Thr232)/Aurora C [1:1000; XP® Rabbit mAb clone D13A11, Cell Signalling), β-tubulin (1:2000; rabbit polyclonal, Abcam ab6046), GAPDH (1:400; rabbit mAb #2118, Cell Signaling Technology). Secondary antibodies used were HRP-conjugated, or IRDye® 680RD- or 800CW-conjugated at 1:10000 dilution for quantitative fluorescence measurements on an Odyssey® Fc Dual-Mode Imaging System (LI-COR Biosciences).

### Ubiquitination Assays

U2OS-bioUb cells transfected with different versions of AURKA-Venus were synchronized as described above and processed for detection of ubiquitin conjugates as previously described (Min 2013).

### Statistical Analysis

Data analyses were performed in GraphPad Prism (San Diego, CA, USA). Results were analyzed with Student’s t-test or Mann-Whitney U-test (non-parametric) as indicated in figure legends. Significant results are indicated as p < 0.05 (*), p ≤ 0.01 (**) or p ≤ 0.001(***). Values are stated as the mean ± standard deviations.

## Data Availability

## Acknowledgements

We thank Norman Davey, Laura Itzhaki and Rohan Eapen for discussions during the course of this work and many past researchers and students in the lab for their contributions to understanding AURKA degradation. This project was supported by Royal Society International Exchanges award (IES/R3/170195) to CL and GG whilst work in CL’s lab was supported by Cancer Research UK (C3/A10239) and BBSRC (BB/R004137/1). Studentship support is acknowledged by AMA from Yousef Jameel Scholarship (Cambridge International Trust), by CNO from Gates Cambridge and Rosetrees Trust, by CA by AstraZeneca UK. AP is funded by Associazione Italiana Ricerca sul Cancro (AIRC MFAG id. 20447).

## Author contributions

CL conceived and wrote the manuscript draft based on ideas developed in discussion with AP and GG. Experiments for the study were contributed by AMA, HBA, CA, CNO, AH, MM, CM, RG, MDL. GJ supervised *in silico* experiments, LA supervised isPLA experiments. CL, AP, GG, GJ, CA, CO reviewed and edited the manuscript.

