## Supplementary Figures for "Revisiting degron motifs in human AURKA required for its targeting by APC/C-FZR1"

### Figure S1

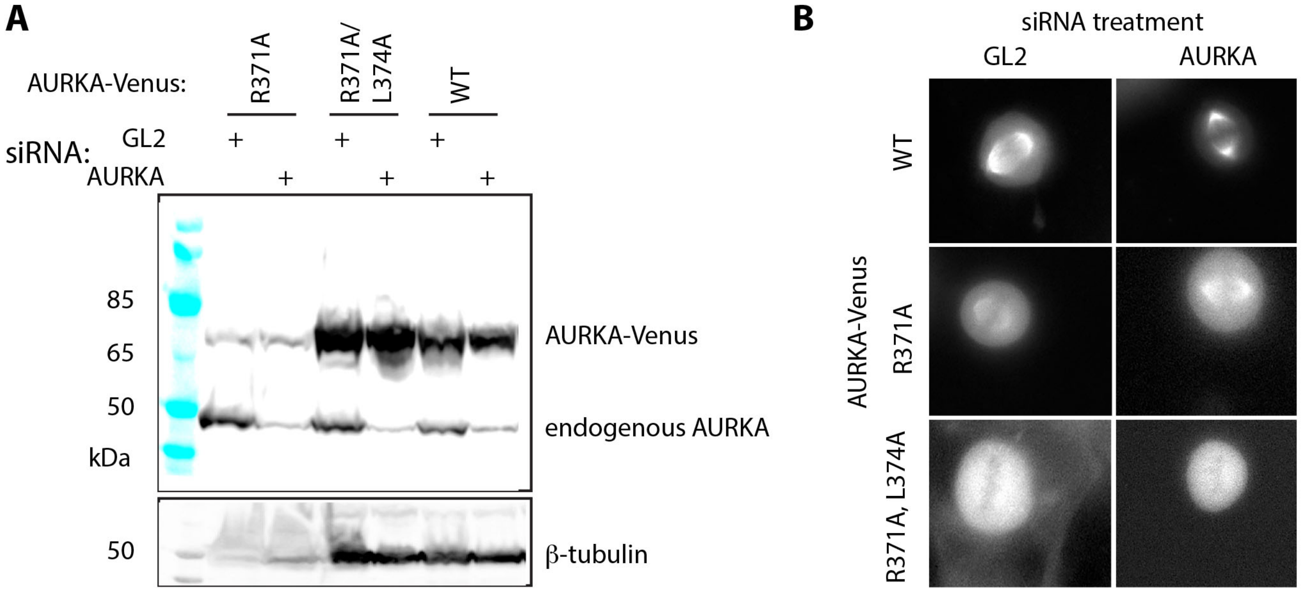


**Localization of C-terminal mutants of AURKA-Venus in absence of endogenous AURKA.** U2OS cells were co-transfected with siRNA-resistant WT or putative D-box mutants of AURKA-Venus and with control (GL2) or AURKA-targeting siRNA. Expression of endogenous and exogenous AURKA was compared by immunoblot (**A**). Example images of mitotic cells captured from fluorescence timelapse imaging (**B**) show that AURKA siRNA does not bring about localization of mutant versions to the mitotic spindle.

### Figure S2

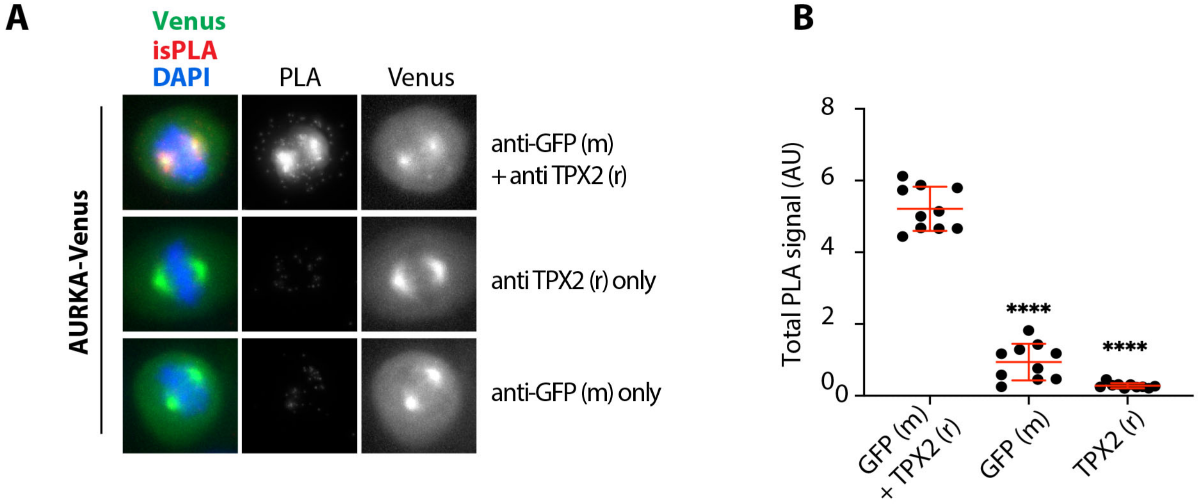


**Controls for isPLA experiment to measure AURKA-TPX2 interaction.** U2OS cells were co-transfected with WT AURKA-Venus and processed for isPLA detection of AURKA-Venus and endogenous TPX2 using mouse GFP and rabbit TPX2 antibodies alone and in combination (**A**). isPLA signal from mitotic cells was quantified over the whole cell (**B**).

### Figure S3

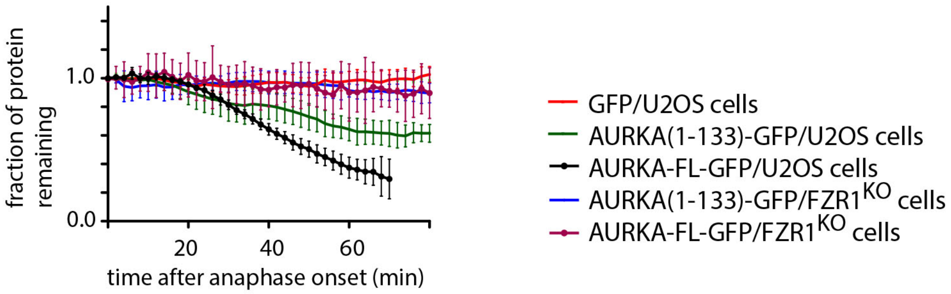


**AURKA(1-133)-GFP is degraded at mitotic exit in FZR1-sensitive manner.**
U2OS parental or FZR1$^{KO}$ cells were transfected with full-length (FL) or 1-133 AURKA tagged with GFP and live cells filmed through mitosis. GFP fluorescence was quantified in individual cells and plotted as fraction of protein remaining relative to level at anaphase onset. Curves show mean values and standard deviations from n≥5 cells. The result was representative of 2 experiments and consistent with Fig 3 data .

### Figure S4

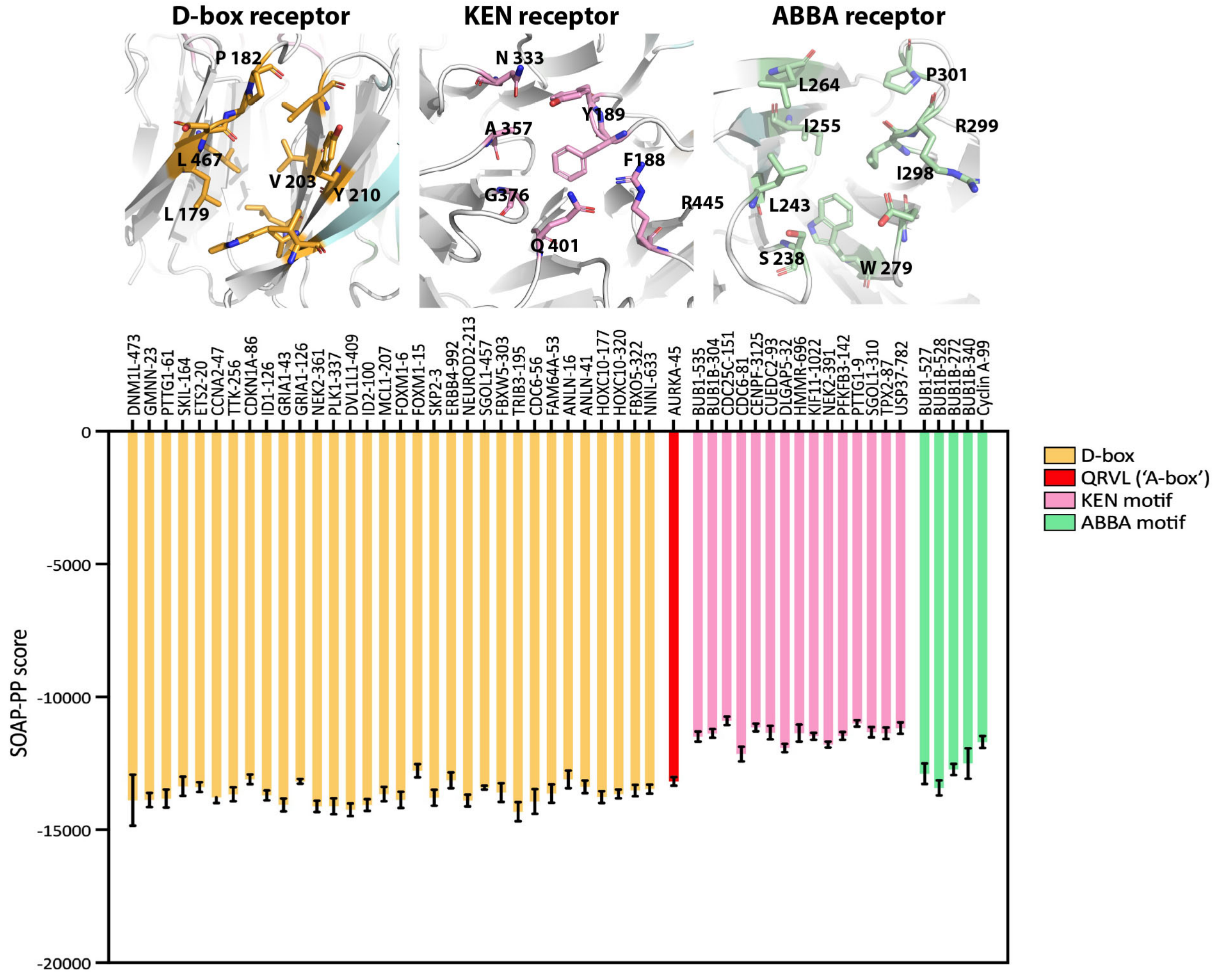


**In silico docking of A-box motif to DBR site compared to a panel of other degrons.**
Peptides corresponding to a range of validated D-box and KEN degrons, the AURKA A-box ($Q_{45}$RVLxxx) and ABBA-motifs that also bind to co-activators (all taken from Davey et al. 2016) were *in silico* docked into the DBR cleft of FZR1. Energies of binding are expressed as SOAP scores and shows that the A-box docks to the DBR as well as many D-boxes.
